# Inhibition of Notch Signaling Attenuates Epileptic Discharges in the Adolescent Rat Brain after Status Epilepticus Induction

**DOI:** 10.1101/2024.11.17.624007

**Authors:** Jie Liu, Jin Chen, Ping Yuan

**Author notes:** **Corresponding author:** Ping Yuan. **Compliance with Ethical Statements**. **Ethical approval:** All protocols were executed according to NIH Guidelines regarding the care and use of laboratory animals and approved and monitored by the Animal Care Committee of Chongqing Medical University, Chongqing, China. **Author contribution :** Ping Yuan and Jin Chen conceived and designed the experiments. Jie Liu and Ping Yuan performed the experiments and analyzed the data. Jin Chen contributed reagents, materials, and analytical tools. Ping Yuan wrote the manuscript.

## Abstract

**Background:** Notch signaling plays a critical role in neuroregeneration after injuries such as those caused by status epilepticus (SE).

**Objective:** To explore the effects of Notch signaling on epileptogenesis and the underlying mechanisms in adolescent rat brains in the acute phase after SE induction.

**Methods:** N-[N-(3,5-difluorophenacetyl)- L-alanyl)]-S-phenylglycine t-butyl ester (DAPT), which indirectly inhibits Notch, was injected into rats during the acute phase after SE induction to inhibit Notch signaling. Electroencephalogram (EEG) was used to observe spontaneous recurrent seizures. Differences in the synaptic structures of the hippocampus were observed by transmission electron microscopy. Nissl staining and Timm staining were used to observe the loss of hippocampal neurons and sprouting of mossy fibers, respectively, in the hippocampus at 28 days after SE.

**Results:** EEG illustrated that DAPT treatment reduced the severity of epileptic discharges after SE induction. Transmission electron microscopy revealed reductions in the presynaptic membrane active band length and postsynaptic membrane dense matter thickness in the CA1 region of the hippocampus. Meanwhile, Nissl staining demonstrated that DAPT treatment reduced the loss of hippocampal neuronal cell degeneration, and the hippocampal structure was repaired to a certain extent. Meanwhile, Timm staining illustrated that DAPT treatment did not affect mossy fiber sprouting (MFS) after SE induction.

**Conclusion:** Inhibiting Notch signaling reduced EEG epileptic activity, attenuated synaptic damage, and partially restored the hippocampal neuronal structure. However, it did not alter MFS after SE induction.

## Introduction

Status epilepticus (SE) is a neurologic emergency describing a prolonged seizure caused by abnormal seizure mechanisms, and this event can lead to neuronal death, injury, and changes in neural networks and cause cognitive dysfunction^1^. SE is accompanied by alterations of the synaptic structure and function within the hippocampus, sustained epileptic discharges, and the onset of mossy fiber sprouting (MFS). During SE, the number of excitatory synapses in the hippocampus increases, whereas the number of inhibitory synapses decreases^2, 3^. Simultaneously, synaptic proteins and receptors involved in neurotransmitter release and synaptic plasticity are altered^4^. In addition, sustained epileptic discharges can propagate rapidly throughout the brain, involving multiple brain regions and disrupting normal neuronal communication, thereby leading to brain energy expenditure and metabolic disruption^5^. Notably, previous studies found that MFS can promote seizures through several mechanisms^6^. Abnormal mossy fibers have been linked to neosynaptogenesis on dentate granule cells, forming recurrent excitatory circuits in the hippocampus. The newly formed connections can bypass normal inhibitory control mechanisms, allowing the balance between excitation and inhibition to be disrupted^7, 8^. This further promotes seizure generation and propagation, and it is believed to play an important role in the development and maintenance of chronic epilepsy^9^. Additionally, hippocampal areas CA1 and CA3 are affected by epileptic activity. It has been suggested that the CA3 region can generate self-sustained epileptiform activity that can propagate to CA1, which can also provide feedback inhibition to CA3 as a regulatory mechanism to limit epileptic propagation^10^. Thus, exploring the range of episodic changes in the hippocampus during sustained epilepsy is crucial for studying the generation and development of epilepsy and its underlying mechanisms.

The Notch signaling pathway plays a crucial role in neuronal development, regulating various processes such as cell fate determination, differentiation, and synaptic plasticity. Notch receptors and ligands are expressed in the developing brain, and their interactions contribute to the proper formation and function of neural circuits^11^. Abnormalities in Notch signaling components have been observed in animal models of epilepsy^12^. The Notch pathway has a pivotal role in epilepsy by influencing the differentiation, proliferation, and dendritic growth and connectivity of neurons^13, 14^. Aberrant Notch signaling can disrupt neuronal differentiation and proliferation and potentially lead to abnormal connections between neurons^14^. Abnormal Notch signaling can affect the balance between excitation and inhibition in neural circuits, leading to hyperexcitability and seizure activity^15^. Additionally, Notch signaling has been implicated in neuroinflammation and gliosis, which contribute to epileptogenesis and the maintenance of chronic seizures^16^. However, the mechanism by which Notch signaling contributes to the development of epilepsy remains unclear.

Our previous research^12^ indicated that the Notch signaling pathway is abnormally activated during the acute phase of SE in hippocampus of rats. In the present study, we investigated the effect of Notch signaling on epileptogenesis in rats after SE induction and explored the potential mechanisms.

## METHODS

### Rat treatment

Sprague-Dawley (SD) rats were fed food and water *ad libitum* and housed in a vivarium under a 12-h/12-h light/dark cycle at 21 ± 1°C and 60% humidity. All protocols were executed according to the NIH Guide for the Care and Use of Laboratory Animals and approved and monitored by the Animal Care Committee of Chongqing Medical University (Chongqing, China). SE was induced in 20-day-old SD rats *via* sequential intraperitoneal (i.p.) injections of lithium chloride (127 mg/kg, Solarbio, Beijing, China) and pilocarpine (30 mg/kg, Sigma-Aldrich, St. Louis, MO, USA). Each animal was randomly assigned to the experimental or control group. Thirty minutes after SE induction, rats in the experimental group were treated with 10 mg/kg (i.p.) diazepam to terminate the seizure, whereas rats in the control group received an injection of the same volume of sterile saline. Atropine l mg/kg was intraperitoneally injected 15 min following SE induction to reduce mortality (20%–30% in this experiment).

After SE induction, rats in the experimental group were randomly assigned to the SE or N-[N-(3,5-difluorophenacetyl)-L-alanyl)]-S-phenylglycine t-butyl ester (DAPT) group. Specifically, 5 μl of DAPT (D5942, Sigma-Aldrich) dissolved in DMSO were injected stereotaxically (0.5 μl/min) into the lateral ventricle of rats in the DAPT treatment group using a Hamilton syringe at 30 min after seizure induction. Conversely, rats in the SE group only received the same volume of DMSO as the vehicle. The experimental design is presented in Figure 1.

**Fig. 1.**
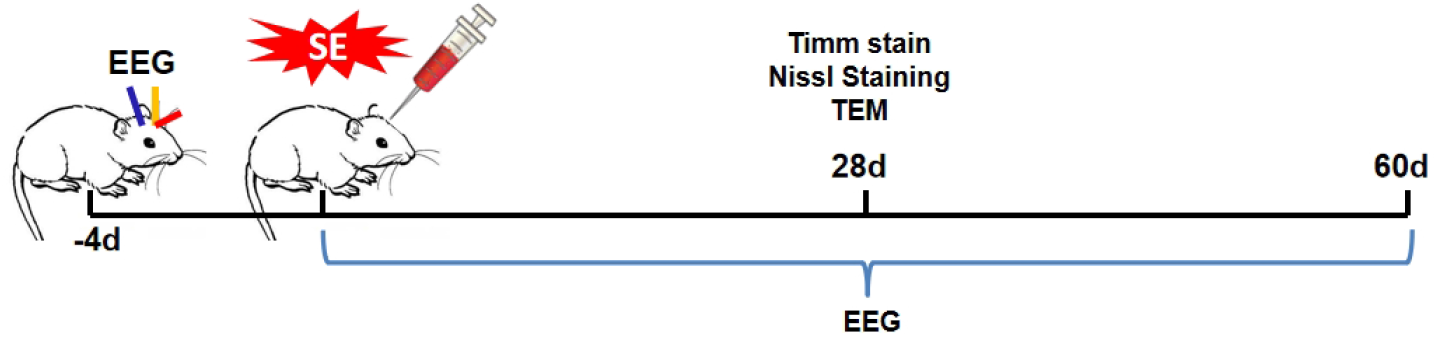
Experimental design. Rats were randomly assigned to the control, SE, or DAPT group. Brain tissues were taken 28 days after SE induction to perform Timm staining, Nissl staining, and TEM. To collect EEG data, rats in the different groups were equipped with electrodes 4 days before SE induction. EEG data were collected continuously for 60 days after SE induction (2 h per day).

### Electroencephalogram (EEG) and analysis

A 1.00-mm-diameter hole was made on the left and right sides of the sagittal suture in the parietal skull of each rat (n = 6/group), and a hole was drilled above the telencephalon for grounding. Then, simple homemade electrodes with 0.1-mm nickel– chromium wires were inserted to touch the dura mater for scalp EEG. After 4 days of recovery, SE was induced to establish the SE model, and EEG was performed during modeling and then for 2 h per day until 60 days after SE induction using a radiophysiological signal detection system (Telemetry ReSCarch Limited, Auckland, New Zealand). EEG was mainly used to record changes in spontaneous recurrent seizures (SRSs). An ADL instrument amplifier was used with the following parameters: low-pass filter at 100 Hz, digitization rate of 2 kHz, and amplification rate of 1000×. According to a previous study^17^, seizures were defined as discharges with the following characteristics: frequency > 5 Hz, amplitude > 2 × baseline, and duration ≥ 10 s. pClampfit software was used to record the average number of daily spontaneous seizures, identify the 10-min period with the most severe seizures, and analyze the frequency of seizures and duration of each seizure in a 10-min period.

### Timm staining to analyze MFS

Timm staining was performed using the FD Rapid TimmStain™ kit according to the manufacturer’s instructions (#PK701, FD NeuroTechnology, Columbia, MD, USA). Specifically, rats were sacrificed 28 days after SE induction (n = 6/group) *via* an intracardial perfusion of sodium sulfide-containing perfusate, followed by 4% paraformaldehyde (PFA) in 0.1 M PBS. Brains were postfixed in 4% PFA for 24 h at 4°C and transferred to 30% sucrose solution in 0.1 M PBS for 72 h at 4°C. Cryoprotected brains were sectioned coronally at 40-μm intervals using a freezing microtome and mounted on positively charged slides, air-dried at room temperature for 24 h, and then stored in a light-protected box at −20°C until staining. Images were acquired using a microscope (Olympus, Tokyo, Japan). MFS was analyzed using a quantitative scale for sprouting in the CA3 and DG regions^18^.

### Nissl staining

CA1 and CA3 neurons from rats in each group were analyzed by Nissl staining 28 days after SE induction. The paraffin sections of brain tissues after different treatments were routinely deparaffinized, dehydrated through an alcohol gradient (soaked in 95%, 80%, and 70% alcohol for 1 min each), and stained by dropping 100–200 μl of Nissl staining solution for 3 min. Distilled water was used to remove the excess staining solution. After dehydration with ethanol and transparentization with xylene, the slices were sealed with neutral gum to complete Nissl staining. Five rats in each group were selected to generate brain tissue specimens, and five sheets of each specimen were selected for Nissl staining. The brain slices on the cover glasses were imaged by fluorescence microscopy (IX53, Olympus). Positive neurons were counted using ImageJ software (US National Institutes of Health, Bethesda, MD, USA).

### Transmission electron microscopy (TEM)

TEM was performed to observe the changes of synaptic morphology and density in the hippocampal CA1 region 28 days after SE induction in the different treatment groups (n = 6/group). Brain tissues were taken after transcardiac perfusion using 4% PFA. Hippocampal CA1 tissues were fixed in 2.5% glutaraldehyde for 2 h at 4°C, washed three times with PBS, and stained with 2% OsO_4_ for 2 h at room temperature. After dehydration *via* a series of alcohol concentrations (30%, 50%, 70%, 80%, 95%, and 100%) and acetone, the tissue was embedded in epoxy resin to form small 5-mm spheres. The embedded tissue blocks were cut into ultrathin sections (60 nm thick) and stained with uranyl acetate and lead citrate. Observation was then performed by TEM.

The criteria used to identify ultrastructural synaptic profiles were previously described^19-21^. To analyze the morphological changes in the length of the presynaptic membrane active band (AD), width of the synaptic gap (SG), and thickness of the postsynaptic density (PSD) in hippocampal synapses at the ultrastructural level, five sections were randomly selected from each sample based on a systematic random sampling principle and photographed by TEM (magnification, ×40,000). The AD length, SG width, and PSD thickness were measured by hand-tracing using ImageJ.

### Statistical analysis

All data were recorded as the mean ± standard error of the mean and statistically analyzed using SPSS 17.0 software (IBM, Armonk, NY, USA). An unpaired Student’s *t*-test was used for comparisons between two groups. For multiple comparisons, one-way analysis of variance with Dunnett’s T3 test was used. *p* < 0.05 was considered statistically significant.

### Data availability statement

All relevant data generated in this research can be requested from the corresponding author.

## RESULTS

### Effect of Notch signaling on epileptic discharges after SE

Epileptogenesis is a pathological process characterized by SRSs that gradually emerge after SE. EEG discharges were recorded daily in the different intervention groups after SE induction, and SRSs of varying degrees, which manifested as scattered sharp waves and spike waves lasting tens of seconds, were detected in all experimental rats in the chronic phase after SE induction (Fig. 2). The duration, frequency, and average number of SRSs per day were further analyzed in each group (Table 1). Frequent epileptiform discharges occurred in the acute phase (1–3 days) after SE induction (Fig. 2b). The number of epileptiform discharges increased slightly over time, but the duration of each epileptiform discharge shortened (Fig. 2c). By contrast, although SRSs also appeared in the chronic period after SE induction in the DAPT group, the amplitude of the sharp and spiky waves decreased, and the frequency of the discharges slowed (Fig. 2d). Based on observation by the naked eye, the behavioral manifestations, which mainly included spontaneous eye gaze and facial muscle twitching and rarely included unilateral or bilateral forelimb clonus, were significantly less severe in the DAPT group than in the SE group. These results suggest that Notch signaling blockade after SE induction partially attenuates the severity of EEG epileptic discharges.

**Table 1.** The SRS duration, seizure frequency, and average number of SRSs per day.

| SRS measurement indicators | SE group | DAPT group | *t*-test | p |
| --- | --- | --- | --- | --- |
| Seizure duration (s/time) | 43.83 ± 8.06 | 33.78 ± 6.86* | 2.326 | 0.0423 |
| Seizure frequency (Hz) | 9.94 ± 3.34 | 5.12 ± 1.62** | 3.181 | 0.0098 |
| Number of SRSs (times/day) | 12.55 ± 4.03 | 8.29 ± 2.27* | 2.256 | 0.0477 |

\* $p < 0.05$ ; \*\* $p < 0.01$ .

**Fig. 2.**
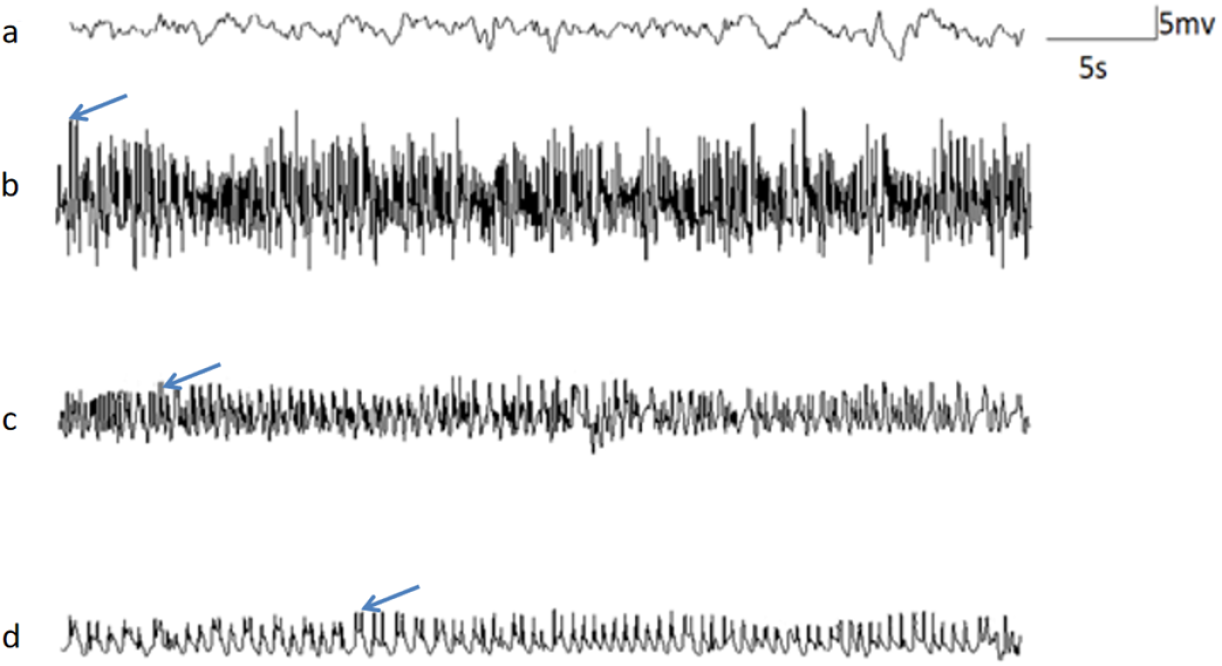
Dynamic changes in EEG after SE induction. A–D: EEG in each group after SE induction (n = 6/group). (a) Control group. (b) SE group. (c) SRSs in the SE group. (d) SRSs in the DAPT group.

### Effect of Notch signaling on the synaptic ultrastructure in the CA1 region of the hippocampus after SE induction

As a vulnerable region in SE, assessing the synaptic ultrastructural changes in the CA1 region of the hippocampus is important for epileptogenesis. Therefore, rats in the SE group were administered DAPT, and the effects of Notch signaling blockade on the synaptic ultrastructure of the CA1 region after SE induction were investigated by TEM (Fig. 3a). The AD length, SG width, and PSD thickness were measured in each group. The results demonstrated that compared with the control group, the AD length was significantly reduced (Fig. 3b), the SG was widened, and the PSD was blurred in the SE group (Fig. 3c). In addition, the thickness of the PSD was reduced (Fig. 3d). By contrast, DAPT administration partially reversed the reductions of the AD length and PSD thickness (Fig. 3b, d), but it had no significant effect on the SG width (Fig. 3c). This suggests that inhibiting the activation of Notch signaling after SE induction can promote the repair of the synaptic ultrastructure.

**Fig. 3.**
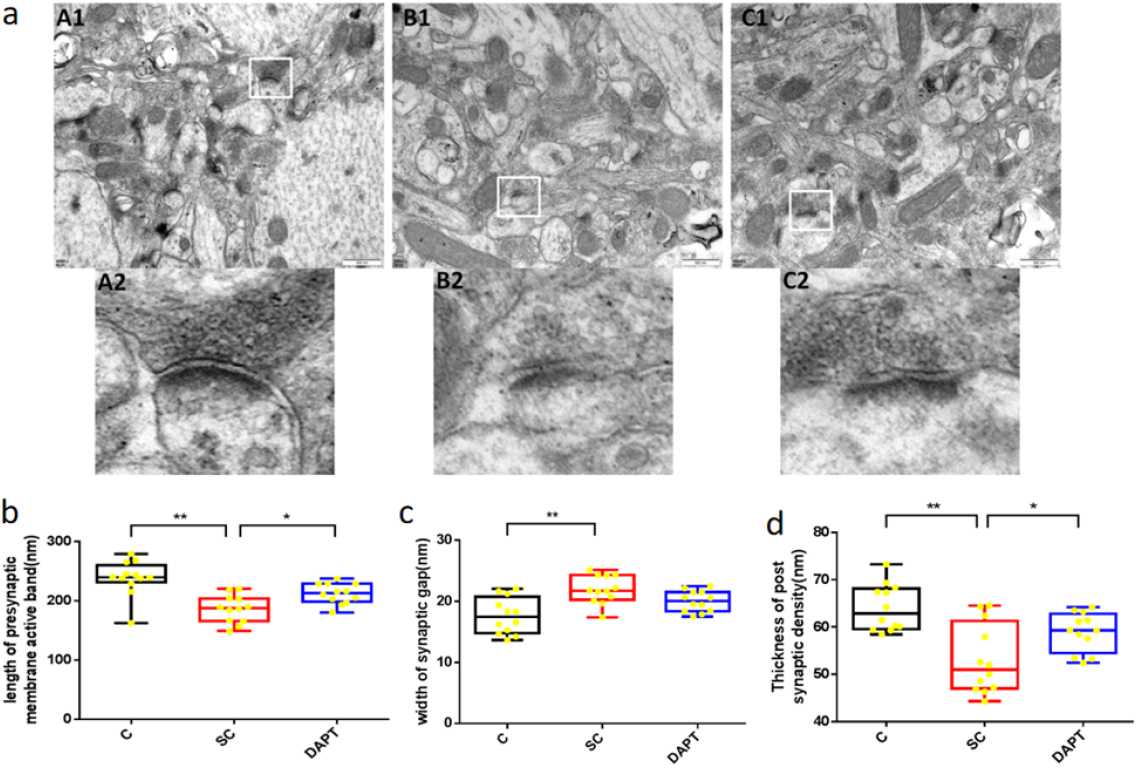
Changes in the hippocampal synaptic ultrastructure after SE induction. (a) TEM images of the hippocampal CA1 region on day 28 in the Control, SE, and DAPT groups. (A1) Control group. (B1) SE group. (C1) DAPT group. Scale bars: 500 nm. b–d: Indicators of the hippocampal synaptic ultrastructure in the different groups. Comparison of the AD length in brain slices (b), SG width (c), and PSD thickness (d) among the control SE, and DAPT groups in the chronic phase.*p < 0.05, **p < 0.01. n = 6/group.

### Effects of Notch signaling on hippocampal neurons after SE induction

Twenty-eight days after SE induction, the hippocampal neuron structure and damage in each group were examined by Nissl staining to determine the effect of Notch signaling on hippocampal neurons. The results illustrated that hippocampal neurons had a loose arrangement in the SE group, and some of the cells exhibited crumpling or vacuolization (Fig. 4a). Notably, compared with the control group findings, there were fewer Nissl bodies in the pyramidal cells of the CA3 and CA1 regions of the hippocampus in the SE group (Fig. 4b), and these findings were accompanied by scattered nucleolysis and disorganized cell arrangement. After DAPT administration, the disorganization of neuronal cells in the CA1 and CA3 regions was significantly improved, and the cellular hierarchy was distinct. The cell size and morphology were basically normalized (Fig. 4a), and the number of homogeneously stained Nissl granules in the cytoplasm was increased (Fig. 4b). Meanwhile, the loss of neurons was alleviated compared with the results in the control and SE groups (Fig. 4a). This suggests that after SE induction, the inhibition of Notch signaling by DAPT partially restored the hippocampal neuronal cell structure and reduced the occurrence of neuronal degeneration and loss.

**Fig. 4.**
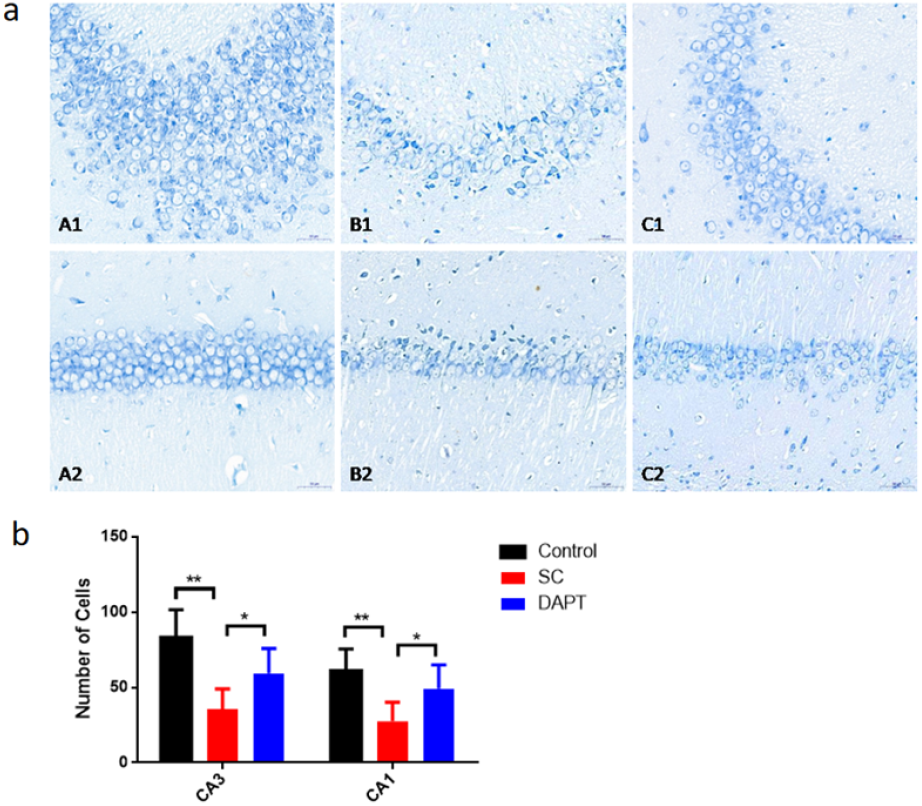
Dynamics of hippocampal neurons after SE induction. (a) Nissl staining of the hippocampal CA3 (A1–C1) and CA1 regions (A2–C2) in the control, SE, and DAPT groups 28 days after SE induction. Scale bar: 50 μm. (b) Comparison of the number of Nissl-stained cells in the CA3 and CA1 regions in the different groups. *p < 0.05, **p < 0.01. n = 5/group.

### Effect of Notch signaling on MFS after SE induction

Timm staining was applied to observe changes in the distribution and number of Timm-stained granules to characterize the distribution of MFS and synaptogenesis. Timm scores and MFS densities were examined on the slices of hippocampal regions from different groups. The MFS density in the hippocampal CA3 and DG regions was significantly increased in the SE group (Fig. 5c), which also exhibited a significantly higher number of Timm-stained particles (Fig. 5a) and higher Timm scores than the control group (Fig. 5b). Notably, the Timm scores and MFS densities in the hippocampal CA3 and DG regions were even higher in the DAPT group, but DAPT treatment did not improve MFS after SE induction (Fig. 5b, c). Thus, our results illustrated that the inhibition of Notch signaling after SE induction did not reduce the occurrence of MFS.

**Fig. 5.**
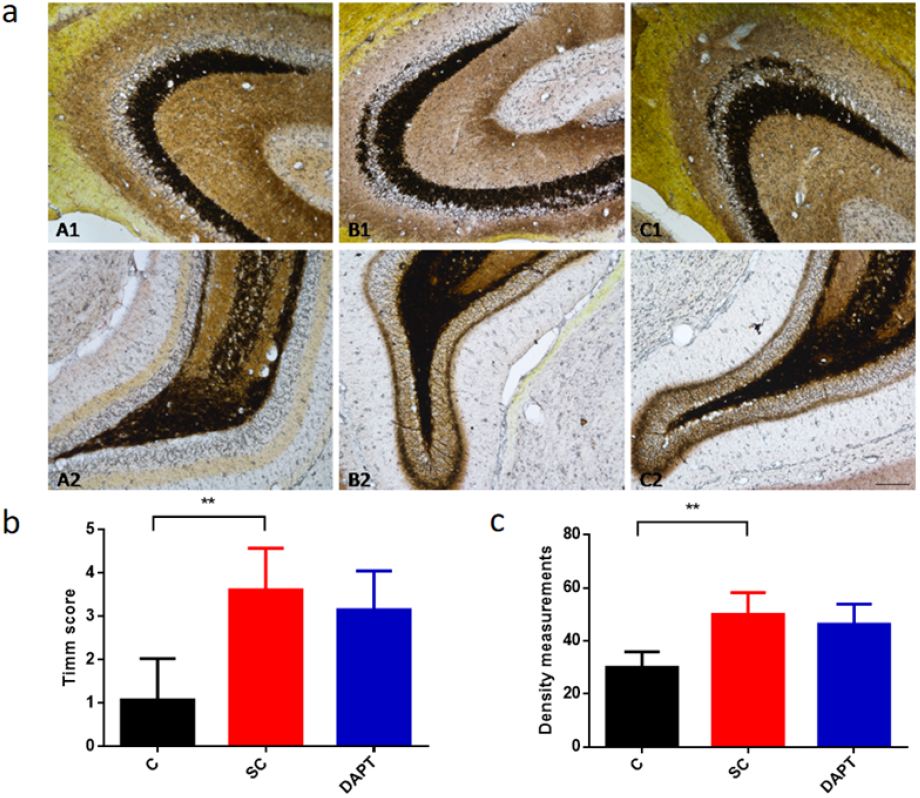
MFS after SE induction. (a) Timm staining of the hippocampal CA3 (A1–C1) and DG regions (A2–C2) in the control, SE, and DAPT groups 28 days after SE induction. Scale bar: 50 μm. (b, c) Hippocampal Timm scores (b) and the MFS density (c) were statistically compared among the treatment groups.*p < 0.05, **p < 0.01. n = 6/group.

## DISCUSSION

We investigated the role of the Notch pathway in epilepsy. Our research revealed that inhibiting Notch signaling with DAPT during the acute phase following seizures effectively reduced the susceptibility to epileptic seizures in the chronic phase in rats, decreasing both the frequency and severity of seizures, in line with previous findings^16, 22, 23^. SRS is a key indicator of epilepsy development after SE development^24^. We also examined SRS development and found that DAPT treatment reduced the severity of epileptic episodes. However, our study also revealed that DAPT treatment did not completely prevent the occurrence of SRSs. All of these findings highlight the complexity of the development of epilepsy as a multifactorial process. Our research illustrated the potential role of the Notch signaling pathway in epilepsy.

The hippocampal neural circuit plays a crucial role in the onset of epilepsy, with abnormal synaptic connections affecting the transmission of information between neurons^16^. This includes both excessive strengthening and weakening of synaptic connections, as well as alterations in pre-synaptic and post-synaptic neurons, leading to instability within the neural network^25^. Previous research suggested that abnormal changes in the synaptic structure can lead to synaptic reorganization, triggering new neural circuits that enhance synchronous firing and excitability, ultimately precipitating epileptic seizures^26^. In recent years, the pivotal role of the Notch signaling pathway in modulating the morphology and function of synapses has received increasing attention^27^. In this study, we utilized DAPT to inhibit Notch signaling, observing notable increases in AD length and PSD thickness. This suggests that inhibiting the Notch signaling pathway contributes to the partial restoration of compromised synaptic structures. However, blocking Notch signaling also underscores the possible negative effects on learning and memory function. This provides new directions and challenges for further research.

The neuronal structure in different hippocampal regions might have distinct roles in different stages of epileptic seizures. For example, once epilepsy occurs, abnormal electrical activity in the CA3 region can rapidly spread to other regions, including CA1, giving rise to the characteristics of generalized epilepsy^28^. In this process, the CA1 region serves as an integrator and transmitter of information, receiving inputs from the CA3 region and other sources and transmitting them to other brain areas, thereby driving the further spread of epilepsy^28, 29^. Additionally, the CA1 region is closely associated with learning and memory^30^, and this region might be disrupted during epileptic states, further highlighting its multifaceted role in epilepsy. The results of Nissl suggested that inhibiting Notch signaling with DAPT after SE induction partially restored the hippocampal neuronal structure and reduced neuronal degeneration and loss in both the CA1 and CA3 regions. These findings align with related research^31-33^, emphasizing the potential role of the Notch signaling pathway in epilepsy development. Contradicting some other studies, our research focused on post-seizure intervention and structural aspects, providing a unique perspective that contributes to a broader understanding of the Notch pathway’s involvement in epilepsy development and highlights the significance of timing and structural considerations in therapeutic approaches. Subsequent studies could unveil the intricate processes involved and strategies to harness their therapeutic benefits.

MFS is one of the most extensively studied pathological mechanisms associated with epilepsy^34, 35^. However, whether MFS is “epileptogenic” or “restorative” remains controversial^6^. Although intrahippocampal circuit reorganization might be a cause of hippocampal epileptiform activity, recent studies indicated that MFS can develop independently of mossy fiber target loss^36^. It is present in animals with spontaneous seizures, but its presence is not necessarily associated with the occurrence of spontaneous seizures^37^. Adult-born granule cells robustly contribute to MFS and form functional recurrent synapses^38^. It has been demonstrated that despite the presence of robust morphological MFS from granule cells born after SE arises, these synapses were not functionally active and unable to drive recurrent excitation^38^. Thus, MFS is neither pro- nor anti-epileptic, and it has also been suggested to be an epiphenomenon^6^. Interestingly, in the present study, inhibition of the Notch signaling pathway by DAPT did not significantly reduce Timm scores and MFS density, suggesting that Notch signaling block after SE does not attenuate chronic phase convulsions by affecting MFS. This discrepancy between our results and previous findings on MFS highlights the complexity of the relationship between Notch signaling and epilepsy^39^.

## CONCLUSION

In summary, this study provided crucial insights into the mechanisms by which the Notch signaling pathway participates in the pathogenesis of epilepsy. By examining various aspects of the impact of this pathway, including its effects on the frequency and severity of seizures, synaptic structure, neuronal structure, and MFS, we obtained a more comprehensive understanding of its potential roles. However, it is essential to acknowledge that the functions of this pathway are diverse, and they might vary because of individual differences. Therefore, future research should seek to increase our understanding of this pathway to facilitate the development of individualized treatments for epilepsy.

## References

1. Trinka E, Cock H, Hesdorffer D, et al. A definition and classification of status epilepticus--Report of the ILAE Task Force on Classification of Status Epilepticus. Epilepsia 2015;56:1515–1523.

2. Andoh M, Ikegaya Y, Koyama R. Synaptic Pruning by Microglia in Epilepsy. J Clin Med 2019; 8:2170.

3. Fan J, Dong X, Tang Y, et al. Preferential pruning of inhibitory synapses by microglia contributes to alteration of the balance between excitatory and inhibitory synapses in the hippocampus in temporal lobe epilepsy. CNS Neurosci Ther 2023; 29:2884–2900.

4. Bourne JN, Harris KM. Coordination of size and number of excitatory and inhibitory synapses results in a balanced structural plasticity along mature hippocampal CA1 dendrites during LTP. Hippocampus 2011;21:354–373.

5. Trivisano M, Specchio N. What are the epileptic encephalopathies. Curr Opin Neurol 2020;33:179–184.

6. Cavarsan CF, Malheiros J, Hamani C, Najm I, Covolan L. Is Mossy Fiber Sprouting a Potential Therapeutic Target for Epilepsy. Front Neurol 2018; 9:1023.

7. Wenzel HJ, Woolley CS, Robbins CA, Schwartzkroin PA. Kainic acid-induced mossy fiber sprouting and synapse formation in the dentate gyrus of rats. Hippocampus 2000; 10:244–60.

8. Luo W, Egger M, Domonkos A, et al. Recurrent rewiring of the adult hippocampal mossy fiber system by a single transcriptional regulator, Id2. Proc Natl Acad Sci U S A 2021; 118:e2108239118.

9. Buckmaster PS, Zhang GF, Yamawaki R. Axon sprouting in a model of temporal lobe epilepsy creates a predominantly excitatory feedback circuit. J Neurosci 2002;22:6650–6658.

10. Huh CY, Amilhon B, Ferguson KA, et al. Excitatory Inputs Determine Phase-Locking Strength and Spike-Timing of CA1 Stratum Oriens/Alveus Parvalbumin and Somatostatin Interneurons during Intrinsically Generated Hippocampal Theta Rhythm. J Neurosci 2016;36:6605–6622.

11. Breunig JJ, Silbereis J, Vaccarino FM, Sestan N, Rakic P. Notch regulates cell fate and dendrite morphology of newborn neurons in the postnatal dentate gyrus. Proc Natl Acad Sci U S A 2007;104:20558–20563.

12. Yuan P, Han W, Xie L, et al. The implications of hippocampal neurogenesis in adolescent rats after status epilepticus: a novel role of notch signaling pathway in regulating epileptogenesis. Cell Tissue Res 2020;380:425–433.

13. Yang Y, He M, Tian X, et al. Transgenic overexpression of furin increases epileptic susceptibility. Cell Death Dis 2018;9:1058.

14. Piacentino ML, Hutchins EJ, Bronner ME. Essential function and targets of BMP signaling during midbrain neural crest delamination. Dev Biol 2021;477:251–261.

15. Hortopan GA, Dinday MT, Baraban SC. Spontaneous seizures and altered gene expression in GABA signaling pathways in a mind bomb mutant zebrafish. J Neurosci 2010;30:13718–13728.

16. Wu L, Li Y, Yu M, Yang F, Tu M, Xu H. Notch Signaling Regulates Microglial Activation and Inflammatory Reactions in a Rat Model of Temporal Lobe Epilepsy. Neurochem Res 2018;43:1269–1282.

17. Cao L, Jiao X, Zuzga DS, et al. VEGF links hippocampal activity with neurogenesis, learning and memory. Nat Genet 2004; 36:827–35.

18. Holmes GL, Sarkisian M, Ben-Ari Y, Chevassus-Au-Louis N. Mossy fiber sprouting after recurrent seizures during early development in rats. J Comp Neurol 1999; 404:537–53.

19. Han W, Pan YN, Han Z, et al. Advanced maternal age impairs synaptic plasticity in offspring rats. Behav Brain Res 2022; 425:113830.

20. Szule JA, Jung JH, McMahan UJ. The structure and function of ‘active zone material’ at synapses. Philos Trans R Soc Lond B Biol Sci 2015; 370:20140189.

21. Gold MG. A frontier in the understanding of synaptic plasticity: solving the structure of the postsynaptic density. Bioessays 2012; 34:599–608.

22. Vezzani A, Maroso M, Balosso S, Sanchez MA, Bartfai T. IL-1 receptor/Toll-like receptor signaling in infection, inflammation, stress and neurodegeneration couples hyperexcitability and seizures. Brain Behav Immun 2011;25:1281–1289.

23. Liu X, Yang Z, Yin Y, Deng X. Increased expression of Notch1 in temporal lobe epilepsy: animal models and clinical evidence. Neural Regen Res 2014;9:526–533.

24. Löscher W, Ferland RJ, Ferraro TN. The relevance of inter- and intrastrain differences in mice and rats and their implications for models of seizures and epilepsy. Epilepsy Behav 2017;73:214–235.

25. Yingjun X, Haiming Y, Mingbang W, et al. Copy number variations independently induce autism spectrum disorder. Biosci Rep 2017;37:BSR20160570 [pii].

26. Engel J Jr, Pitkänen A. Biomarkers for epileptogenesis and its treatment. Neuropharmacology 2020;167:107735.

27. Billiard F, Lobry C, Darrasse-Jèze G, et al. Dll4-Notch signaling in Flt3-independent dendritic cell development and autoimmunity in mice. J Exp Med 2012;209:1011–1028.

28. Kim SR. Control of Granule Cell Dispersion by Natural Materials Such as Eugenol and Naringin: A Potential Therapeutic Strategy Against Temporal Lobe Epilepsy. J Med Food 2016;19:730–736.

29. Palacio S, Chevaleyre V, Brann DH, Murray KD, Piskorowski RA, Trimmer JS. Heterogeneity in Kv2 Channel Expression Shapes Action Potential Characteristics and Firing Patterns in CA1 versus CA2 Hippocampal Pyramidal Neurons. eNeuro 2017;4:ENEURO.0267-0217.2017.

30. Wu Z, Li X, Zhang Y, Tong D, Wang L, Zhao P. Effects of Sevoflurane Exposure During Mid-Pregnancy on Learning and Memory in Offspring Rats: Beneficial Effects of Maternal Exercise. Front Cell Neurosci 2018;12:122.

31. Yoon KJ, Lee HR, Jo YS, et al. Mind bomb-1 is an essential modulator of long-term memory and synaptic plasticity via the Notch signaling pathway. Mol Brain 2012;5:40.

32. Zhang ZJ, Guo MX, Xing Y. [ERK activation effects on GABA secretion inhibition induced by SDF-1 in hippocampal neurons of rats]. Zhongguo Ying Yong Sheng Li Xue Za Zhi 2015;31:443–447.

33. Liu C, Ying Z, Li Z, et al. Danzhi Xiaoyao Powder Promotes Neuronal Regeneration by Downregulating Notch Signaling Pathway in the Treatment of Generalized Anxiety Disorder. Front Pharmacol 2021;12:772576.

34. Song MY, Tian FF, Wang YZ, Huang X, Guo JL, Ding DX. Potential roles of the RGMa-FAK-Ras pathway in hippocampal mossy fiber sprouting in the pentylenetetrazole kindling model. Mol Med Rep 2015;11:1738–1744.

35. Wang X, Yu Y, Ma R, Shao N, Meng H. Lacosamide modulates collapsin response mediator protein 2 and inhibits mossy fiber sprouting after kainic acid-induced status epilepticus. Neuroreport 2018;29:1384–1390.

36. Ad R, Santhakumar V, Howard A, Soltesz I. Mossy cells in epilepsy: rigor mortis or vigor mortis. Trends Neurosci 2002; 25:140–4.

37. Nissinen J, Lukasiuk K, Pitkänen A. Is mossy fiber sprouting present at the time of the first spontaneous seizures in rat experimental temporal lobe epilepsy. Hippocampus 2001; 11:299–310.

38. Hendricks WD, Chen Y, Bensen AL, Westbrook GL, Schnell E. Short-Term Depression of Sprouted Mossy Fiber Synapses from Adult-Born Granule Cells. J Neurosci 2017; 37:5722–5735.

39. Song Y, Wang C, Cai H, et al. Functional hierarchy of the angular gyrus and its underlying genetic architecture. Hum Brain Mapp 2023;44

